# Biocompatible hydrogel electrodeposition enables the simultaneous preparation of multi-microinterfaces for ligand bioconjugation and multiplexed electrochemical detection

**DOI:** 10.1101/2023.08.17.553669

**Authors:** Vuslat B. Juska, Nadia Moukri, Pedro Estrela, Graeme D. Maxwell, Stacey Hendriks, Esmeralda Santillan-Urquiza, Bernadette O’Brien, Bernardo Patella, Rosalinda Inguanta, Alan O’Riordan

## Abstract

Affinity based electrochemical biosensing systems with integrated miniaturised interfaces has enabled key advancement toward rapid, sensitive, precise and deployable detection platforms. Manufacturing silicon micro/nano devices for biology interface has been a highly promising platform to design and develop electrochemical sensors for the detection of very low concentrations of the target molecules. However, the biofouling challenge of the biosensors when the surface is exposed to a complex matrix such as blood, serum, milk, has been a road blocker. Here we introduce a simple, rapid formation of an anti-biofouling coating onto several electroactive surface areas present on a single chip simultaneously. Using such a multiplexed surface, we were able to investigate the optimum working conditions on-chip. Concentrating on two individual bioassay platforms for stress biomarkers, haptoglobin and cortisol, we demonstrate the broad applicability of the developed universal platform with excellent performance in bovine serum and correlation with conventional ELISA using milk samples.

## 1. Introduction

Electrochemical sensing based bioassay development has been a fast growing research field due to the benefits of the overall platform such as miniaturisation, multiplexing, and rapid response. Silicon microtechnology is a highly effective way of achieving the fabrication of the miniaturised chips with design flexibility allowing to pattern metal surfaces at micro and nanoscale with multiple active areas on a single device^1-2^. On the other hand, such technology provides the opportunity not only to miniaturise the classical three-electrode based electrochemical cell but also to scale down the electroactive surfaces to ultramicro and nanometers with multiplexing capabilities^3-6^. Most electrochemical systems use the well-known external reference electrodes, e.g., Ag/AgCl and saturated calomel electrode (SCE)^7-12^; but, when the reference electrode material is a metal such as gold or platinum, then it is important to explore the working conditions of the current system due to the possible potential shift^6, 13^. Another significant consideration is the fact that the electrical output of these devices is dependent on the electroactive surface properties, therefore, achieving a clean surface is vital for the construction of the sensor^6^. Further, convenient surface functionalization is important for reliable device response after the sensing surfaces are exposed to highly complex matrices such as blood, saliva, serum, milk or other media.^14^ Because the non-target biomolecules present in the sample may attach to the surface, which then provides a false positive response. Such drawback has an impact on the biosensor performance and commercialisation of the affinity based sensing systems^15^. For example, the home-use glucometer strips are the excellent examples for electrochemical biosensors, with a very successful commercialisation history, have been able to overcome this limitation by using an initial blood filtering step to remove the large biomolecules when the sample touches the probe^16-17^. This minimises the interaction of the other biomolecules with the detection surface. Nevertheless, the size-dependant-separation approach is impractical for immunosensors, because the size of the target biomolecules is approximately the same as of other non-target biomolecules present in biological fluids. Therefore, the success of an immunosensor relies on the surface properties of the designed sensing system.

To achieve reliable immunosensing systems and boost the performance of affinity-based sensors, several design strategies have been studied for the development of anti-biofouling surfaces. The most relevant methods are polyethylene glycol (PEG), hydroxy-functional methacrylates (HEMA, HPMA), zwitterionic polymers or monoethylene glycolated (MEG) ^18-21^ based surface modifications. However, these methods have their different problems such as rapid oxidation^22^, expensive, complex and time-consuming synthesis. Alternatively, Bovine Serum Albumin (BSA) integrated immobilisation surfaces have demonstrated very promising results to minimize non-specific bindings. BSA is typically included in the blocking step of various biological assays such as biosensors^23-26^, blots^27^ and enzyme-linked immunosorbent assays (ELISA)^28^. Recent studies have shown the advantage of combining the BSA molecules into the immobilisation matrix for immunosensing. For example, Kim et al.^29^ demonstrated the immobilisation of anti-cortisol antibodies onto a thermally denatured BSA protein layer. To further improve the anti-biofouling properties of BSA based surface matrixes, Sabaté Del Río et al.^30^ demonstrated the use of a gel matrix consisting of glutaraldehyde crosslinked BSA and gold nanowires. The developed antifouling gel was applied to a clean gold surface *via* overnight incubation and the resulting interface was resistant to biofouling up to one month. Li et al.^31-32^ also developed two individual protocols; i.e., in the first approach, they treated BSA with Tris(2-carboxyethyl)phosphine hydrochloride (TCEP) to prepare amyloid-like BSA which was then crosslinked with conductive polymer polyaniline (PANI); in the second approach they electrodeposited the PANI nanowires on the electrode surface and then treated with BSA and glutaraldehyde for crosslinking step

Building upon our previous studies, electrodeposition is one of the best ways to achieve selective, fast and one-step depositions onto macro^14, 33^ or micro/nanoscale electroactive areas^6, 8^. Here, we present the simultaneous one-step electrodeposition of biocompatible antifouling layers consisting of chitosan, BSA and graphene oxide onto microelectrodes with disk shapes. Following this we demonstrate the miniaturisation of the electroactive surface as well as the three-electrode setup into a compact silicon chip. The fabricated chip is suitable to be used for varying electrochemical setup; e.g. (1) the developed device can be used in a conventional three electrode setup by dipping the device in a electrochemical cell for electropolymerisation, (2) the device can be used with hydrogen bubble template due to the adjusted distance of the sensing electrodes with connection pads, (3) the device can be used for drop-casting for surface modification, and (4) the device can be used with ca. 10 μL volume of redox solution covering the three sensing electrodes, reference and counter electrodes for on-chip measurement. The fabrication protocol includes photolithography, deposition, lift-off and etching. After the fabrication, we have characterised the device with electrochemical and microscopic techniques. We used the fabricated chips for the application of miniaturised hydrogen bubble template in order to achieve the deposition of gold foam (AuFoam) nanostructures since the porous AuFoam increases the electroactive area dramatically^6-8^. AuFoam surfaces were used to explore the electro-co-deposition of denatured BSA *via* a biocompatible hydrogel chitosan in the presence of nanostructures for electrical enhancement such as graphene oxide. To achieve a universal site selective immobilisation of IgG isomer antibodies, we first immobilised Protein A/G onto the antifouling coating *via* glutaraldehyde. Then, the several IgG isomers such as anti-Haptoglobin and anti-Cortisol antibodies were linked to the Protein A/G *via* their heavy chain, F_c_. We have shown the efficiency of the developed immunosensing surface without the need for a secondary antibody in serum samples. We have demonstrated the response of the sensors on chip in buffer as well as in a complex matrix such as bovine serum. Resulting immunoassays were studied with milk samples collected from cows and results compared with ELISA as a gold standard method.

## 2. Experimental Section

### 2.1. Materials

Unless otherwise mentioned, all chemicals were used as received. Chitosan, Bovine Serum Albumin (BSA), reduced graphene oxide, acetic acid, potassium chloride (KCl), sodium chloride (NaCl), potassium ferrocyanide (K_4_[Fe(CN)_6_]), potassium ferricyanide (K_3_[Fe(CN)_6_]), acetone, sterile human serum, phosphate buffer saline tablets (PBS, 0.01 M, pH7.4), gold(III)chloride trihydrate, sulphuric acid, glutaraldehyde were obtained from Sigma-Aldrich. Protein A/G, IgG binding buffer, Superblock, anti-Cortisol and anti-Mouse IgG(Alexa Fluor Plus 405) were purchased from Thermofisher. Anti-EGFR (Alexa Fluor 405) was purchased from ABCAM. Fetal Bovine Serum (FBS) was obtained from Generon. Graphene oxide water dispersion was obtained from Graphenea. Anti-Haptoglobin and Bovine Haptoglobin protein were obtained from Fitzgerald (*via* 2BScientific).

### 2.2. Microfabrication, hydrogen bubble templating and antifouling coating

We have previously reported several fabrication protocols of silicon based gold devices by using the major microfabrication techniques including photolithography, deposition, etching and lift-off^6-8, 34-35^. Building upon this experience, we have designed two individual silicon devices; e.g. device 1 is individual disk shaped working electrode with 180 μm diameter and the device 2 is a multiplexed silicon devices with three sensing electrodes (100 μm diameter each) and one pair of reference/counter electrodes. Device 1 is dedicated for antifouling assessment with external reference/counter electrodes and Device 2 is dedicated for sensing application on-chip without a need for external electrodes.

#### Device Microfabrication

Fabrication of the devices is described in previous reports applying different set of masks, varying number of lithography steps and the different thicknesses for the materials depisition^6-8, 35^. In here, a silicon wafer is processed for the thermal growth of silicon dioxide layer with a thickness of 1.5 μm. Each device had two sets of masks; e.g. metal mask and passivation mask. First the metal photolithography was applied in order to achieve the pattern transfer of the devices and following this the lift-off is applied to remove the excess gold. Then a silicon nitrate passivation layer is applied. Finally, sensing electrodes of both devices, reference and counter electrodes of device 2 and connection pads are etched. Before mechanical dicing, wafers were coated with a protective layer of a resist. Fig. 1a demonstrates the representative image of fabrication flow for Device 1 “sensing electrode” with 180 μm diameter disk shaped sensing electrode area.

**Figure 1.**
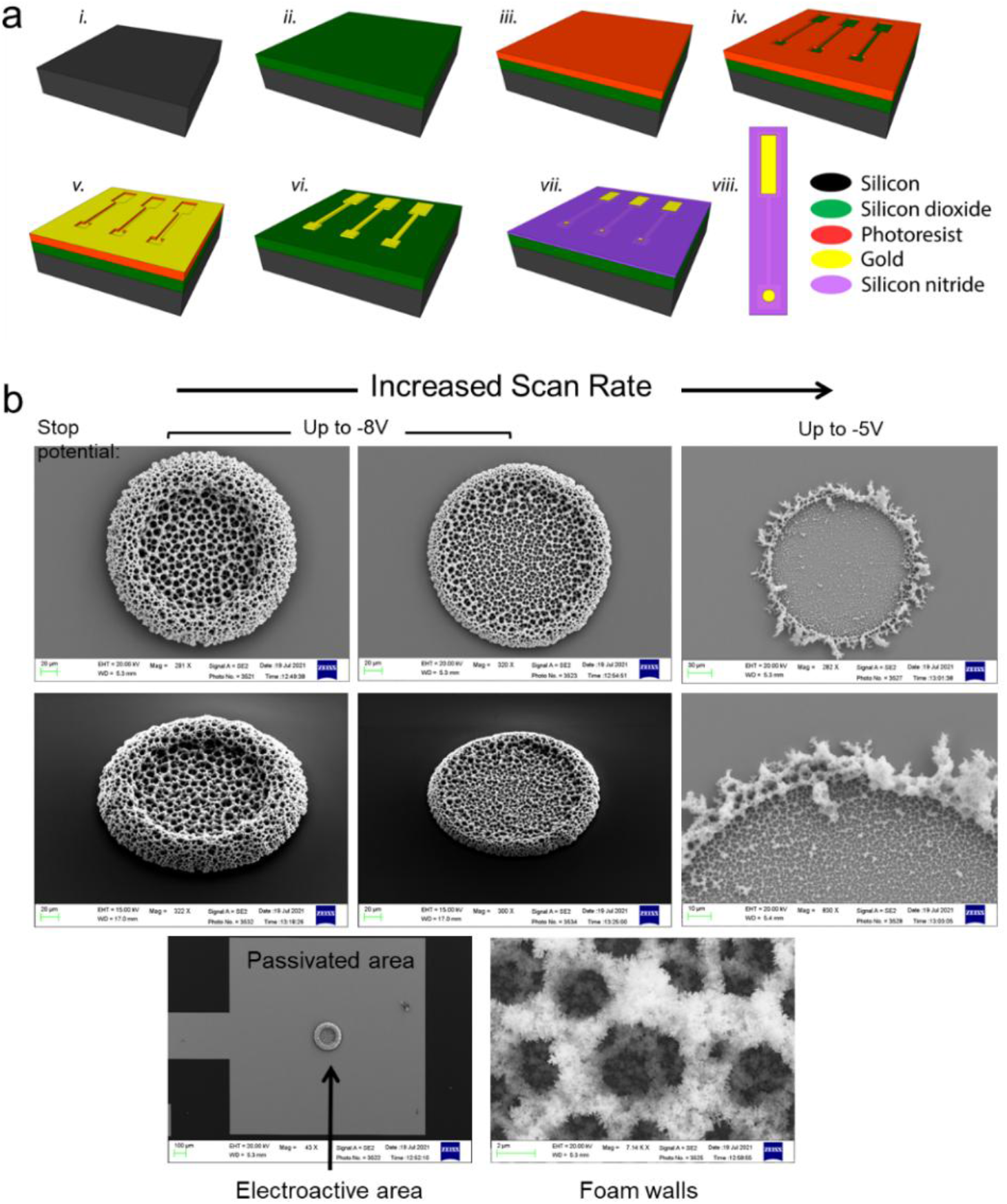
Summary of fabrication and characterisation of Aufoam deposition of Device 1, (a) schematic image of the fabrication flow (I. silicon wafer, ii. oxide growth, iii. Photoresist, iv. Pattern transfer, v. Ti:Au deposition, vi. Lift-off,, vii. Passivation coating), (b) Characterisation of the Aufoam deposition protocol with applied potentials up to -5 V and -8V with increased scan rates.

#### Hydrogen Bubble template for AuFoam deposition

After fabrication, devices washed with acetone and deionized water several times. Then we have modified them with gold foam (AuFoam) depositions *via* a hydrogen bubble template, similarly our previous studies^7-8^. We have optimised the deposition conditions with scan rate and applied stop voltage of cyclic voltammetry as shown in Fig. 1b. The hydrogen bubble template studied by applying a cyclic voltammetry in a solution of 2.8 M H_2_SO_4_ contained 48 mg HAuCl_4_·3H_2_O. For the devices prepared for antifouling assessment study, the scan rate was kept at 100 mV s^-1^ during the CV based gold foam deposition protocol and the highest applied negative voltage was -8 V in order to achieve the highest surface area.

#### Antifouling gel coating

The electrodeposition solution of the antifouling coating was prepared freshly just before use. Chitosan solution was prepared in a glass bottle as previously described^8^, e.g. 0.1 g chitosan flakes were dissolved in 20 mL 0.1 M acetic acid solution. The solution mixed approximately for 3 hours and then stored at room temperature. Thermally denatured BSA was prepared freshly before use. For this, 5 % BSA solution was treated with 5 cycles of 50 °C for 5 minutes and 70 °C for 2 minutes in a thermocycler. Briefly, into a beaker 5850 μL deionized water, 50 μL of graphene oxide, 100 μL BSA and 200 μL chitosan were added subsequently and after each addition the solution mixed very well. The antifouling coating depositions were achieved *via* electrodeposition using cyclic voltammetry.

### 2.3. On-chip immunosensor

#### Sensor preparation

The antifouling gel coated chips were washed with deionized water and excessive 0.2 M phosphate buffer (pH 7.0) subsequently. The chips dried at room temperature. 1.5 % glutaraldehyde solution was freshly prepared before use by diluting 25 % glutaraldehyde (grade I) in 25 mM phosphate buffer (pH 7.0). The 5 μL of 1.5 % glutaraldehyde was drop-casted onto three sensing electrode area. The chips were placed in a petri dish saturated with water vapour and kept in the fridge (+ 4 °C) over night. Following this, the chips were washed with excessive phosphate buffer (25 mM, pH 7.0) and dried at room temperature. The sensor surface was coated with 5 μL of 200 ng/mL Protein A/G solution (25 mM phosphate buffer, pH 7.0). The chips were incubated for two hours at room temperature. Following this, chips were washed gently with excessive phosphate buffer in order to remove weakly attached Protein A/G molecules. The surface was blocked with 5 μL of Superblock® for 20 minutes. After the blocking, the chips were washed to remove weakly attached blocking molecules. The resulting Protein A/G surface was incubated with 40 μg/mL anti-Haptoglobin and/or anti-Cortisol antibodies (IgG binding buffer®) for 75 minutes at room temperature. The resulting immunosensor chips were stored at +4 °C until used. For calibration study, the target molecule in PBS (0.7 μL per electrode) was dropped onto electrodes and incubated for 40 minutes at room temperature. The complex matrix analysis was studied with the haptoglobin or cortisol spiked in whole fetal bovine serum samples and incubated for 40 minutes.

Electrochemical measurements were studied with differential pulse voltammetry (DPV) using a potentiostat (PGSTAT302N, Autolab Metrohm, UK). DPV signals were obtained with a potential step of 5 mV, pulse amplitude 50 mV, pulse modulation time of 50 ms and a pulse period of time 100 ms. The antibody immobilised surfaces (background of the immunoassay) were obtained by measuring DPV until the equilibrium allowing a repeatable DPV response. The peak height was calculated using Nova 2.1.5 software. Signal changes that corresponded to target protein attachment were calculated with the target response subtracted from background.

### 2.4. Milk collection, ELISA and immunosensor

Milk samples were collected from cows at Teagasc Research dairy farm (Moorepark, Ireland). A 70 mL aliquot of pooled milk was collected aseptically from individual cows at morning milking. Teat ends were disinfected with 70 % isopropyl alcohol wipes and the first 3 strips of milk discarded from each quarter before collection. All samples were immediately placed on ice and then stored at –80 °C until analysed.

Milk samples were analysed for cortisol and haptoglobin at Teagasc Food Research Centre. Enzyme linked immunosorbent assay (ELISA) was chosen to determine the concentrations of cortisol and haptoglobin as a gold standard method; e.g. Bovine Haptoglobin Sandwich ELISA Kit (Catalogue No. orb549212; Biorbyt Ltd. Cambridge, UK) and Cortisol ELISA Kit (catalogue No. OKEH02541; Aviva Systems Biology. California, USA). Each ELISA was deployed following the manufacturers’ instructions. The analytical range of the haptoglobin assay reported by the manufacturer is 156 to 10,000 ng/mL, with a sensitivity of 41.5 ng/mL. The analytical range of the cortisol assay reported by the manufacturer is 1.56 to 600 ng/mL, with a sensitivity of <0.31 ng/mL. Absorbance measurements for Haptoglobin, and Cortisol were read at 450 nm using an automatic plate reader (Model Synergy HT; Bio-Tek Instruments, Inc., VT). Standard curves for Haptoglobin and Cortisol were obtained using a 4-parameter logistics (4PL). Unknown sample concentrations were interpolated from the linear portion of the standard curve and results analysed and calculated using Prism (version 9.1.0; GraphPad software; San Diego, CA).

### 2.5. Scanning Electron Microscopy (SEM) and Confocal Laser Scanning Microscopy (CLSM)

SEM imaging was carried out on a Carl Zeiss Supra 40 Scanning Electron microscope that provides high resolution imaging at high and low voltage in high vacuum: 1.3nm @15 kV. The microscope was equipped with SE detector, In-lens detector, Backscatter detector and STEM detector.

cLSM imaging was carried out on a confocal laser-scanning microscope (FluoView 1000, Olympus) equipped with 60X (PLAPON, NA1.35) and 100X (UPLSAPO, NA1.4) oil immersion objectives. The microscope was equipped with three filter cubes mounted on the scope; DAPI/Hoechst (ex 360-370, em 420-500), FITC/EGFP/Bodipy/Fluo3/Di O(ex 470-490, em 515-555) and TRITC, Rhodamine, Di I (ex 530-550, em 590-640).

## 3. Results and Discussion

### 3.1. Anti-biofouling coating design and development

We have assessed the antifouling properties of the coatings by carrying out cyclic voltammograms in a redox solution before and after overnight incubation in buffer containing 1 % BSA^30^. For this study, the best approach is first to assess the antifouling properties of the coating on a slightly larger electrode surface and then miniaturise the coating. Therefore, we have used the Device 1 with 180 μm diameter disk shaped electrodes and we further applied a hydrogen bubble template to increase the overall electroactive surface area of the gold electrode due to the formation of highly porous gold foam deposits.

We first tested the anti-biofouling properties of gold foam (AuFoam) and gold foam/chitosan-reduced graphene oxide (AuFoam/CS-rGO) modified electrodes. While AuFoam showed the same current density after overnight 1% BSA incubation, the peak-to-peak distance (ΔE_p_) of AuFoam increased drastically indicating limited diffusion of ferri/ferrocyanide to the electrode surface due to the clogged pores of the AuFoam (Fig.2a). When we measured the antifouling properties of AuFoam/CS-rGO, we observed a decrease on the current, and ΔE_p_ (Fig.2a) of the coated surface. Clearly, CS-rGO coating allows the accumulation of BSA molecules on the surface resulting in the apparent decrease on the current. In order to overcome this limitation of the coating, we added heat treated BSA (0.008 %, 0.012 %, 0.016 %, 0.08 %, 0.12 % and 0.16 %) to the coating composition (Fig.2b). The best anti-biofouling property is obtained with the 0.08 % BSA co-deposited CS-rGO coating, which provided a 4 % increased current and decreased ΔE_p_ after overnight incubation with 1 % BSA solution.

**Figure 2.**
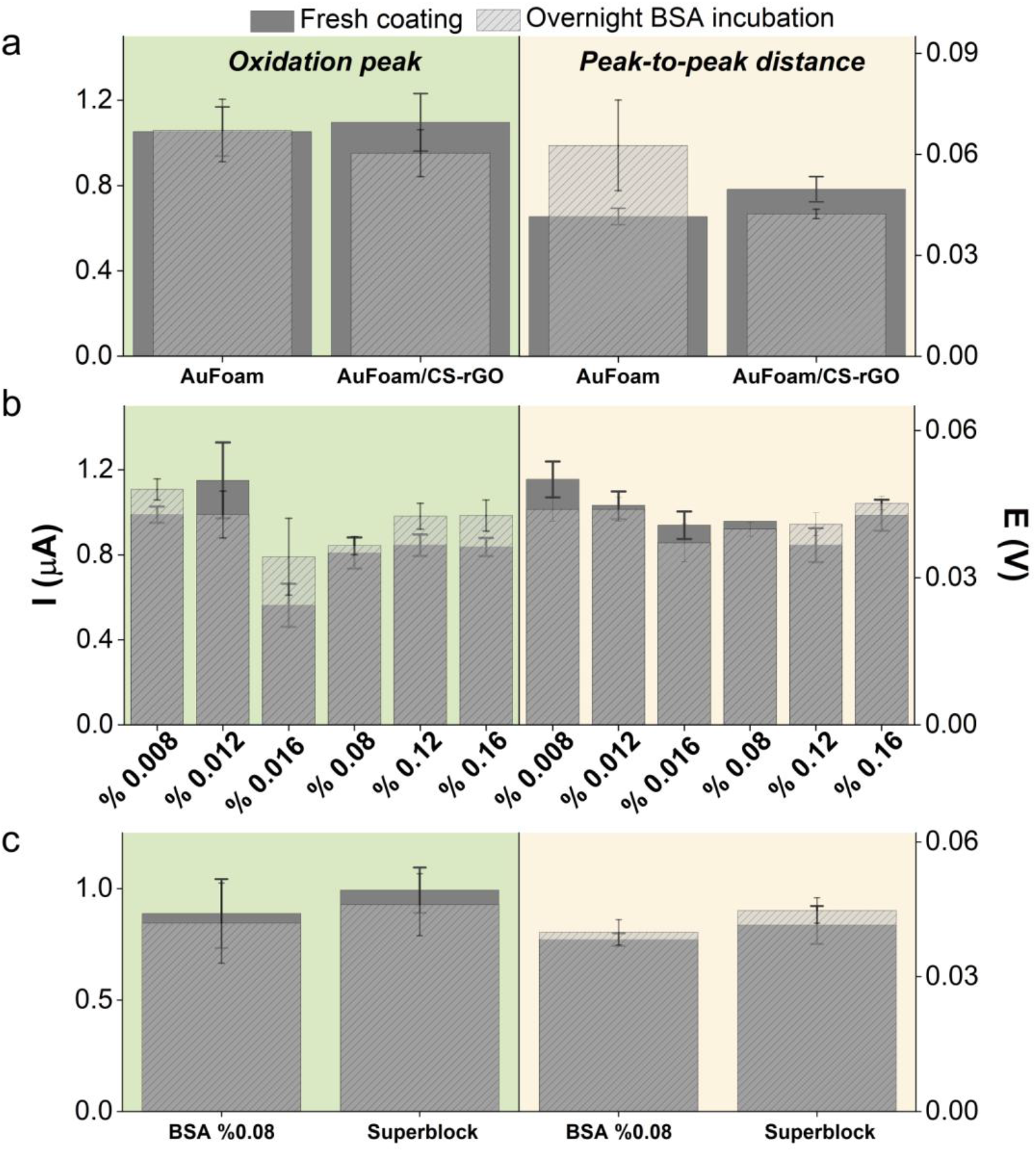
Anti-biofouling assesment. (a) Oxidation peak and Peak-to-peak distance comparison of AuFoam and AuFoam/CS-rGO before and after incubation with 1 % BSA (overnight). (b) Comparison of the surfaces in the presence of varying concentrations of BSA (0.008, 0.012, 0.016, 0.08, 0.12 and 0.16 % BSA). (c) Anti-biofouling capability comparion of BSA and Superblock (ThermoFisher).

Then, we assessed the anti-biofouling properties of AuFoam/CS-rGO-BSA (0.08%) coating with sterile serum. For this, we incubated the coated electrodes with sterile serum overnight. The developed coating showed a 5 % decreased current and 4 % increased ΔE_p_ (Fig.2c), thus reflecting the overall success of the anti-biofouling coating. We further assessed the anti-biofouling properties of Superblock (a commercial product of ThermoFisher). To prepare the coating, we electro-co-deposited Superblock with CS-rGO. The developed AuFoam/CS-rGO-Superblock coating showed 7 % decreased current (Fig.2c) and 8 % increased ΔE_p_ (Fig.2c), thus indicating the good antifouling properties of the coating based on Superblock.

### 3.2. Miniaturisation and multiplexing: Device 2

We have designed and microfabricated a compact silicon chip (Device 2). The chip has three sensing electrodes, one pair of reference and counter electrodes. To achieve the designed device, we employed silicon microtechnologies which provide high reproducibility over the resulting chips^36^. The microfabrication has two steps of lithography - one for pattern transfer, and the other for passivation (Fig.3a). This technology allows us to mass-produce silicon based chips. We produced >111 chip copies per 100 mm diameter silicon substrate (Fig.3b). The overall surface area of the chip is 7.8 × 5.8 mm^2^. The varying application protocols were considered for the design of the chip, e.g. the distance of the sensing electrodes is 1.48 mm allowing the individual drop casting by using a lab scale regular micropipette, and the distance between the connection pads and the sensing electrodes is 4.66 mm. Therefore, the design allows the easy and simultaneous deposition of the foam structures onto three sensing electrodes. Figure 3c shows the voltammetry applied for the deposition of the Au foam and SEM images of the surfaces from different voltage steps. We determined the maximum negative applied voltage (Lower vertex potential) to be -4 V since it provided enhanced surface area with high reproducibility as it could be seen in Fig. 3d,e.

**Figure 3.**
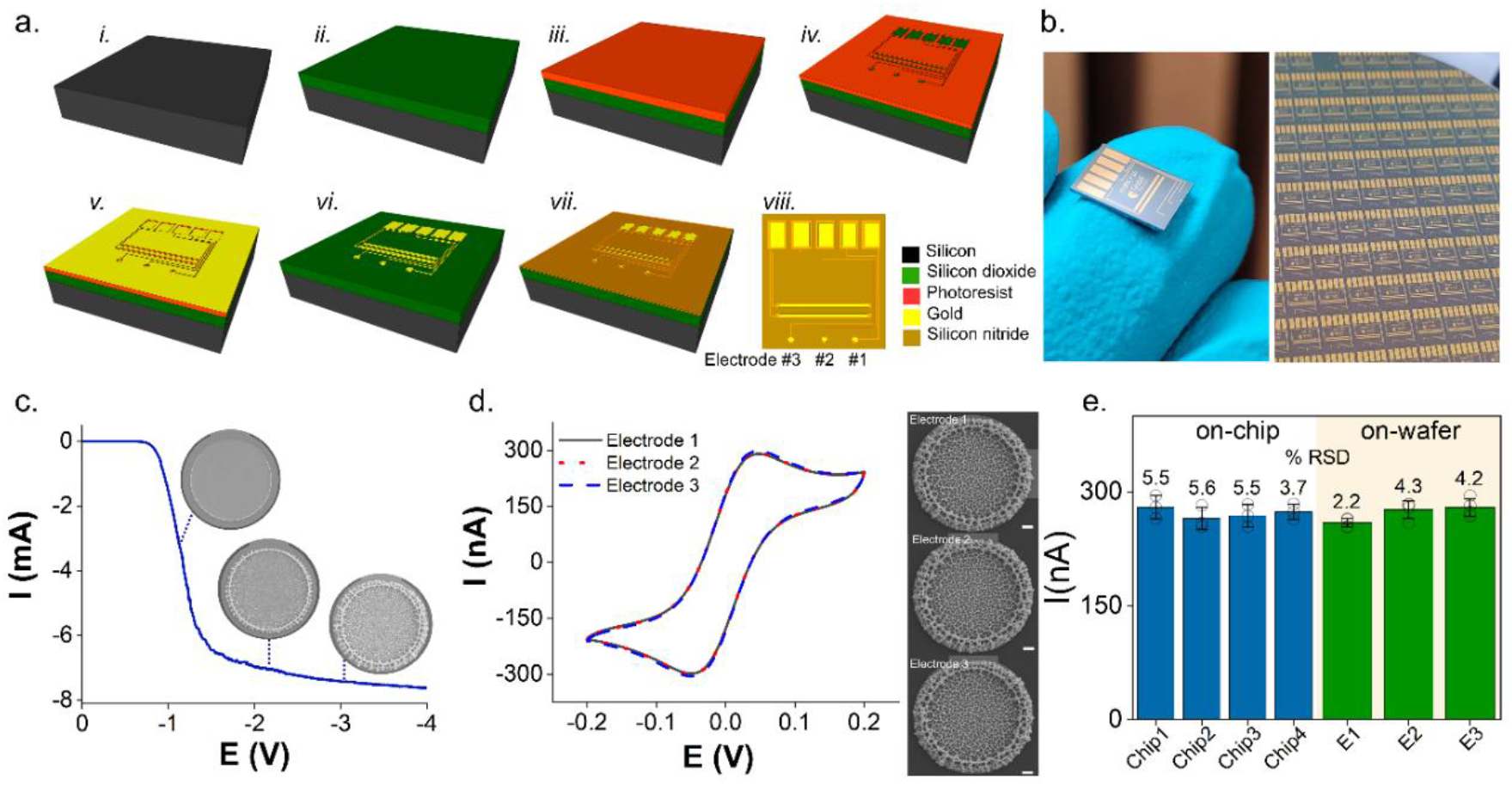
Multiplexed device, Device 2. (a) Microfabrication flow (i. Si wafer, ii. Oxide growth, iii. Photoresist, iv. Pattern transfer-lithography, v. Ti:Au deposition, vi. Lift-off, vii. Passivation, viii. Top image of chip showing the three electrodes, reference and counter electrodes, connection tracks and pads). (b) Photograph of a single chip on a finger tip and the whole wafer. (c) AuFoam deposition protocol via CV. SEM images shows the step by step foamtion of the AuFoam. (d) Overlapping voltammograms of each senisng area after Aufoam deposition and the corresponsding SEM images. (e) On-chip and on-wafer reproducbility of the devices after Aufoam deposition.

### 3.3. Functionalization and surface characteristics via microscopy

The Au foam deposited multiplexed chip was immersed in the antibiofouling solution (i.e. chitosan/BSA/graphene oxide) for the application of the 2^nd^ electrochemical deposition *via* cyclic voltammetry^8, 14^. Fig. 4a summarizes the steps followed for the electro-co-deposition simultaneously onto each sensing area on a single chip. In the absence of graphene oxide we observed a dramatic decrase on the oxidation and reduction current peaks of the chitosan/BSA layer (Fig. 4b), which was then improved with the addition of graphene oxide allowing an efficient electron transfer (Fig. 4c). On the other hand, the high concentrations of the nanoparticles embedded into hydrogels may clog the pores of the matrix after polymerisation that has previously been demontrated with alginate and copper nanoparticles^37^. Therefore, we have studied the several concentrations of the graphene oxide nanosheets in the deposition solution, in order to improve the electrical properties of the miniaturised antifouling surface area. Clearly, 0.05 mg graphene oxide exhibited the most conductivity and ideal matrix condition. The resulting antifouling layers were studied with scanning electron microscope (Fig. 4d) to observe the applied coating. The SEM images show the homogenously distributed thin layer of the antibiofouling film of biocompatible gel.

**Figure 4.**
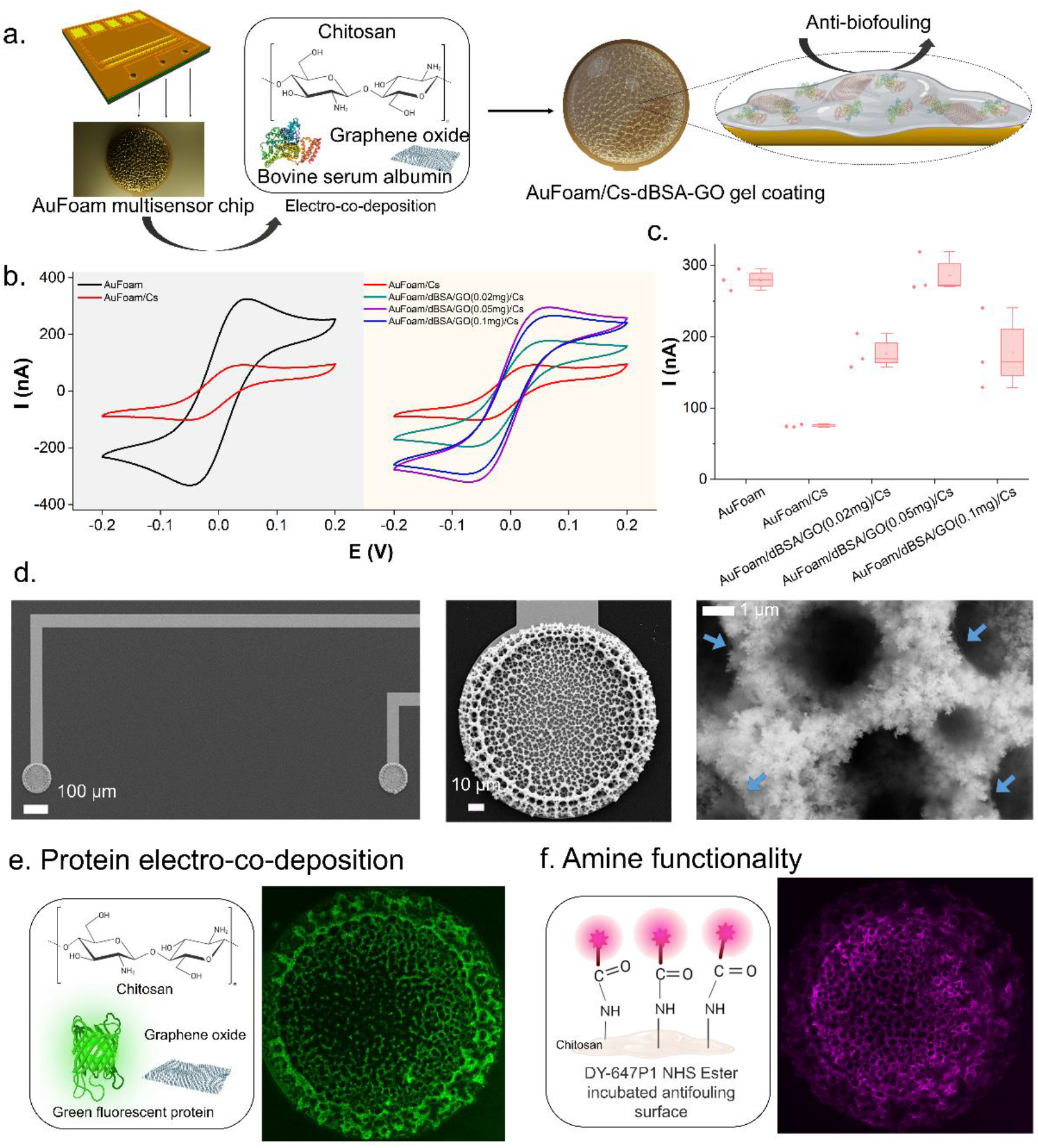
Anti-biofouling surface. (a) Multiplexed chip with three sensing area modified with AuFoams. This surface is applied electro-co-deposition protocol in order to achieve simultaneous deposition of the anti-biofouling coating onto three senisng electrodes. (b) Voltammograms obtained in a redox solution of 5mM FeCN ^3-/4-^ of several surface compositions inclusing AuFoam, AuFoam/CS, AuFoam/ dBSA/GO(0.02 or 0.05 or 0.1 mg)/CS. (d) Graph showing the reproducibility of studied surfaces. (d) SEM images of the Anti-biofouling coating applied onto AuFoam. Blue arrows indicate the thin film of anti-biofouling. CLSM images of (e) electro-co-deposited GFP and (f) DY-647P1 (NHS-ester) modified anti-biofouling surface.

We further investigated the biofunctionilty of the coating *via* confocal laser scanning microscopy (CLSM). First we assessed the success of the electro-co-deopisition of a protein *via* chitosan electropolymerisation. For this, we replaced BSA in the antifouling deposition solution with green fluoresent protein (GFP)^38-39^. Then the GFP electro-co-deposited chitosan surface was invetigated with a confocal microscope^40-41^. Excitation of GFP was performed with a 488 nm Multiline Argon Laser. Fig. 4e shows the Confocal microscopic analysis of the GFP co-depositied surface. A homogenously distibuted fluorescence intensity of GFP was observed and that data visiually demonstrates the success and efficiency of the electro-co-deposition of the proteins *via* a chitosan electropolymerisation step. Second, we assesed the amine functionalilty of the antibiofouling coating since chitosan has abundant amine groups that is one of the advantage of this polymer allowing the bioconjugation of the molecules *via* crosslinking with high yield^8, 42-46^. In order to demonstrate the existence of the surface amines of the antifouling layer, we have incubated the chips with NHS ester DYE (DY647P1) with an excitation wavelenght of 647 nm (the applied excitation was 633 nm *via* a Red Helium Neon Laser). Since the dye had activated carboxyle groups *via* carbodiimide chemistry, these groups can easily react with the secondary amines available on the surface, resulting in amide bonds. The distribution of the fluoresence intensity was homogenous over the foam surface, demonstrating the abundant amine groups (Fig.4f).

### 3.4. Crosslinking and bioconjugation

Our approach for the bioconjugation of the biomolecules was the directed immobilisation of the antibodies in order to enhance the biosensing performance of the devices since random immobilisation protocols do not guarantee the correct immobilisation form of antibodies^47^. Motivated by the literature examples on oriented immobilisation protocols of antibodies^48-51^, we sought to investigate whether a recombinant Protein A/G could be an effective way to attach a wide range of IgG antibodies onto 100 μm disk sensing areas on-chip and thus established a universal platform^52^. Protein A/G is a recombinant protein with the immunoglobulin Fc binding domains from both Protein A (four domains) and Protein G (two domains), providing high affinity towards all IgG species and subclasses recognised by either Protein A or Protein G. Therefore, Protein A/G can be considered as an excellent capture protein for a wide range of IgG antibodies. We achieved the attachment of Protein A/G onto anti-biofouling layer by crosslinking *via* glutaraldehyde chemistry and then we applied two different IgG isomer antibodies, i.e. anti-Haptoglobin and anti-Cortisol for haptoglobin and cortisol detection from milk samples. Fig. 5a summarizes the bioconjugation protocol and applied electrochemical method.

**Figure 5.**
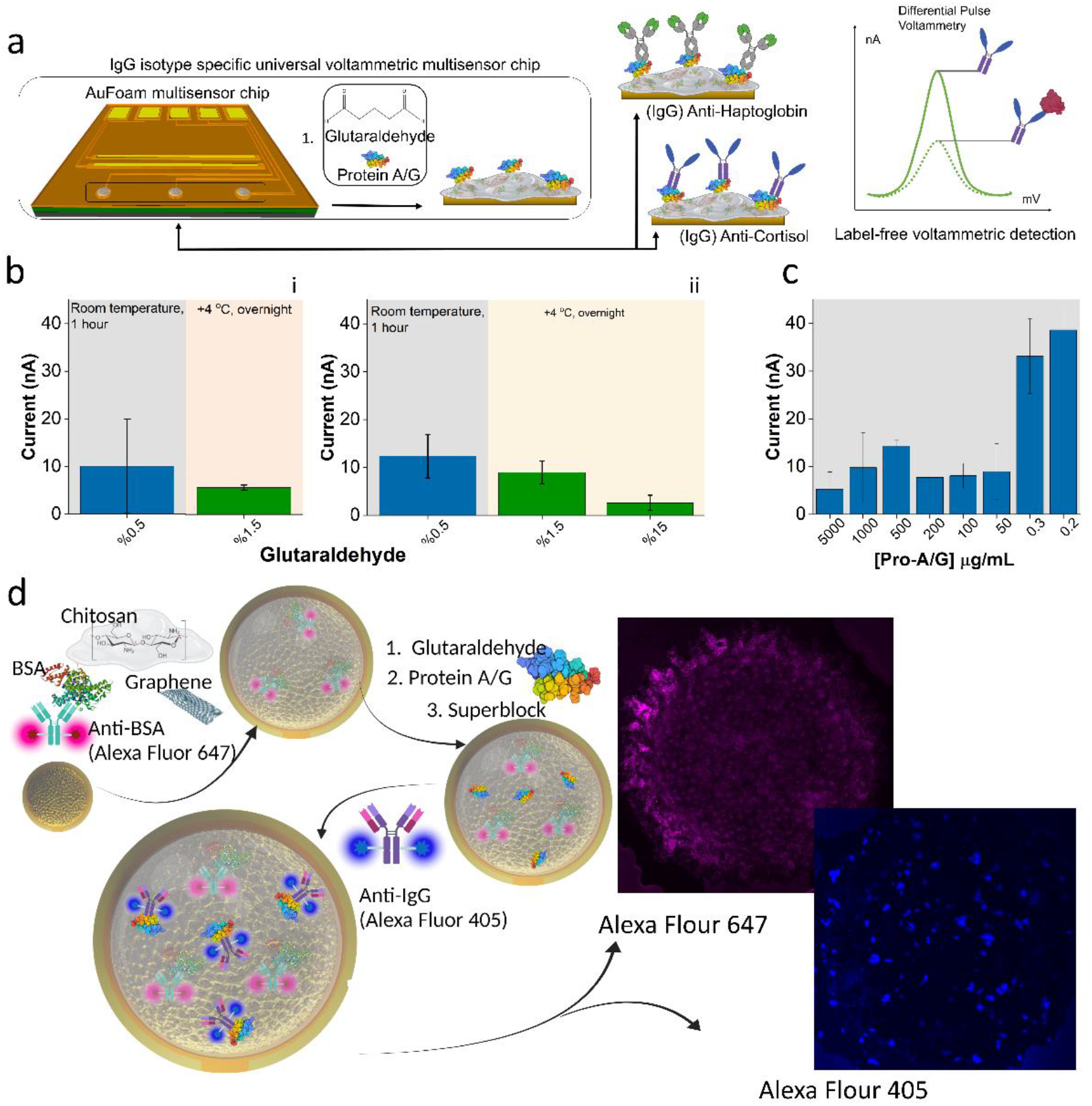
Electrochemical bioassay. (a) Preparation of the bioassays onto Aufoam/anti-biofouling deposited 100-μm disk shaped electroactive surfaces via glutaraldehyde activation and Protein A/G attachment. This follows the incubation of the anti-IgGs (anti-Haptoglobin and anti-Cortisol). Detection technique is DPV. Measurement is taken after antibody incubation (background peak) and then same surface is measured after target incubation (response peak). (b) Glutaraldehyde activation; (i) the graph demonstrates the results of the haptoglobin sensing platform constructed with 5mg/mL Protein A/G, and activated with 0.5 % (1 hour) and 1.5 % (overnight) glutaraldehyde, (ii) the graph demonstrates the haptoglobin sensing platform constructed with 0.1 mg/mL Protein A/G, and activated with 0.5 %(1 hour) and 1.5 % - 15 % (overnight) glutaraldehyde. (c) Optimisation of the Protein A/G concentration which was studied with a wide range from 5mg/mL to 200 ng/mL Protein A/G. (d) Confocal scanning microscopy sample preparation protocol: The schematic image shows the electro-co-deposition of the BSA in the presence of anti-BSA(tagged with Alexa Fluor 647)-and graphene. Then the glutaraldehyde activation and protein A/G immobilisation and superblock incubation. Following this affinity reaction between the anti-IgG (tagged with Alexa Fluor 405) and immobilised protein A/G. CLSM images of Alexa Flour 647 (pink) and and Alexa Flour 405 (blue) were taken using the same sample.

A variety of bioconjugation conditions was studied in order to determine the ideal glutaraldehyde activation protocol and the protein A/G concentration for the surfaces. First, we tested the surfaces incubated with 25 mM phosphate buffer containing 0.5 % glutaraldehyde for monomeric activation at room temperature for 1 hour ^53^. Then the surface was incubated with 5 mg/mL Protein A/G, Superblock and anti-Haptoglobin antibodies respectively. The anti-Haptoglobin based sensor surface was used to target haptoglobin protein present in PBS solution; however, in this protocol of glutaraldehyde activation, the response of the developed bioassay did not provide high reproducibility, which can be attributed to the moderate rate of glutaraldehyde activation for coupling amines of Protein A/G. Following this, we increased the glutaraldehyde concentration to 1.5 % and incubated overnight at + 4 °C, which may result high reactivity for amine coupling *via* glutaraldehyde activation (Fig. 5b, i). Despite the decreased response of the bioassay towards the target haptoglobin protein, we obtained a high reproducibility with this activation protocol. In order to amplify the sensor response we decreased the applied Protein A/G concentration because the incubation with the high concentration may result in the denaturation of the immobilised molecules due to the conformational change of three-dimensional structure^34^. To test this hypothesis, we incubated anti-biofouling surfaces with 0.5 % glutaraldehyde at room temperature and then we immobilised 0.1 mg/mL Protein A/G (50 times decreased concentration). This activation protocol enhanced the bioassay response and reproducibility as expected (Fig. 5b,ii). Further improving the assay conditions, we assessed the 0.1 mg/mL Protein A/G concentration with 1.5 % and 15 % glutaraldehyde (incubation conditions: overnight at + 4 °C). Clearly, we observed an increase on the response when we incubate with 1.5 % glutaraldehyde overnight. In order to achieve a dimeric activation protocol, which may result in a high reactivity towards amine groups, we also tested 15 % glutaraldehyde incubated surfaces overnight; however, we observed a dramatic decrease on the response. Therefore, we did not use the higher concentrations of glutaraldehyde for activations protocol as we met our most reproducible and improved response with 1.5 % glutaraldehyde concentration. Since we witnessed an enhanced biosensor response with a decreased concentration of Protein A/G for anti-Haptoglobin immobilisation, we assessed a wide range of concentration of protein A/G starting from 5000 μg mL^-1^ to 0.2 μg mL^-1^. We observed that the high concentrations of protein A/G didn’t provide very sensitive sensors when the surfaces were used for anti-Haptoglobin immobilisation to detect Haptoglobin; however, a significant increase on the biosensor response was achieved when the Protein A/G concentration was in 0.2 μg/mL. Clearly, the Protein A/G concentration and glutaraldehyde activation play key importance in the development of effective sensing mechanism in here.

Following this, we investigated the characterisation of the bioconjugation protocol *via* confocal laser scanning microscopy (CLSM) by developing an immunofluorescence based surface (Fig. 5d). Briefly, we electrodeposited the anti-biofouling surface in the presence of anti-BSA tagged with Alexa Flour 647. Then, we activated the surface with glutaraldehyde (1.5 %, overnight at + 4 °C) and immobilised Protein A/G (0.2 μg/mL). We applied Superblock for 20 mins and incubated the Protein A/G immobilised surface with anti-IgG tagged with Alexa Flour 405. The excitation of Alexa Flour 405 was performed with a 405 nm UV Diode Laser and the excitation of Alexa Flour 647 was performed with a 633 nm Red Helium Neon Laser (Fig.5d). Having observed electro-co-deposition of GFP *via* chitosan electropolymerisation previously, we therefore also demonstrated the existence of BSA in the presence of anti-BSA (tagged Alexa 647). Using confocal microscopy, the homogenously distributed BSA and protein A/G-anti-IgG (tagged Alexa 405) were shown^54^.

### 3.5. Electrochemical bioassay performance

The developed approach was adapted to construct two affinity-based electrochemical bioassays by immobilising the anti-Haptoglobin and anti-Cortisol antibodies onto Protein A/G functionalised anti-biofouling matrix. A label free voltammetric assay protocol based on DPV was carried out for each sensing system. We first assessed the analytical performance of the bioassays toward the target molecules prepared in buffer for the calibration of each sensing system (Fig. 6 a, b). The sensitivity of the assays was investigated in parallel and both were capable of detecting very low concentrations of targets; 10 fg mL^-1^ of cortisol and 18 fg mL^-1^ for Haptoglobin. This very low detection limits can be attributed to the ultra-small electroactive surface area and the site selective immobilisation protocol *via* Protein A/G. Cortisol assay demonstrated a dynamic range from 10 fg mL^-1^ to 5 pg mL^-1^ (Fig. 6a) and the Haptoglobin assay showed a dynamic range from 18 fg mL^-1^ to 4.8 pg mL^-1^ (Fig. 6b).

**Figure 6.**
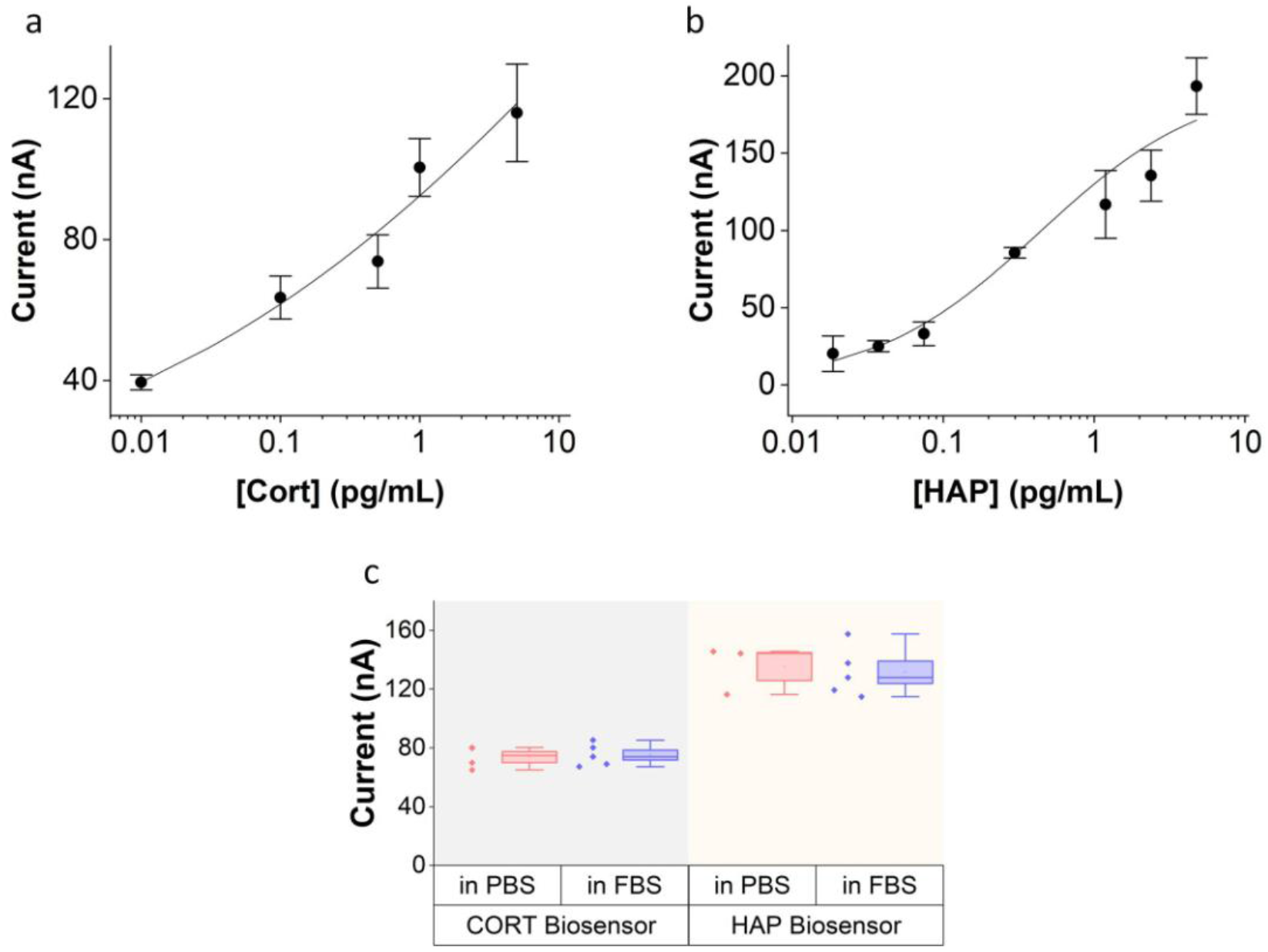
Electrochemical bioassay performance (n≥ 3 for each data set). (a) Cortisol bioassay calibration curve and (b) haptoglobin bioassay calibration curve. Chips were exposed to PBS with spiked target molecules. Hill fitting was performed to fit the calibration data in both assays (adjusted R2 was 0.952 and 0.972 for cortisol and haptoglobin, respectively.) (c) Complex matrix study in FBS and comparison with results obtained in PBS; target concentrations: 0.5 pg/mL cortisol and 2.38 pg/mL haptoglobin.

Once we established the analytical properties of the bioassays, we pursued to investigate their use in complex matrixes. For this purpose, the bioassays were performed with the fetal bovine serum (FBS) samples, which were spiked with the cortisol and/or haptoglobin. Fig. 6c confirms the response of the cortisol and haptoglobin assay in PBS and FBS containing 0.5 pg mL ^-1^ cortisol and 2.38 pg mL^-1^ haptoglobin, respectively. As it can be seen, the assays demonstrated excellent performance when they exposed to a whole complex matrix.

### 3.6 Milk analysis

To further validate the developed multiplexed anti-biofouling universal electrochemical bioassay platform, we analysed eight milk samples obtained from cows. The obtained milk samples were first studied with ELISA to determine the cortisol and haptoglobin concentrations Prior to ELISA, the milk samples were centrifuged at 10,000 *g* for 10 minutes at room temperature to separate milk into a skimmed milk. Samples were diluted and investigated with ELISA following the instructions.

The samples were then studied with the on-chip biosensing platform. For on-chip sensing, we did not apply a centrifugation; however, in order to meet the criteria of our biosensors’ calibration range, we diluted the milk samples in PBS. Diluted milk samples were drop-casted on to the sensor surface and incubated for 40 mins. The sensors results were measured in parallel with the ELISA values as seen in Fig. 7.

**Figure 7.**
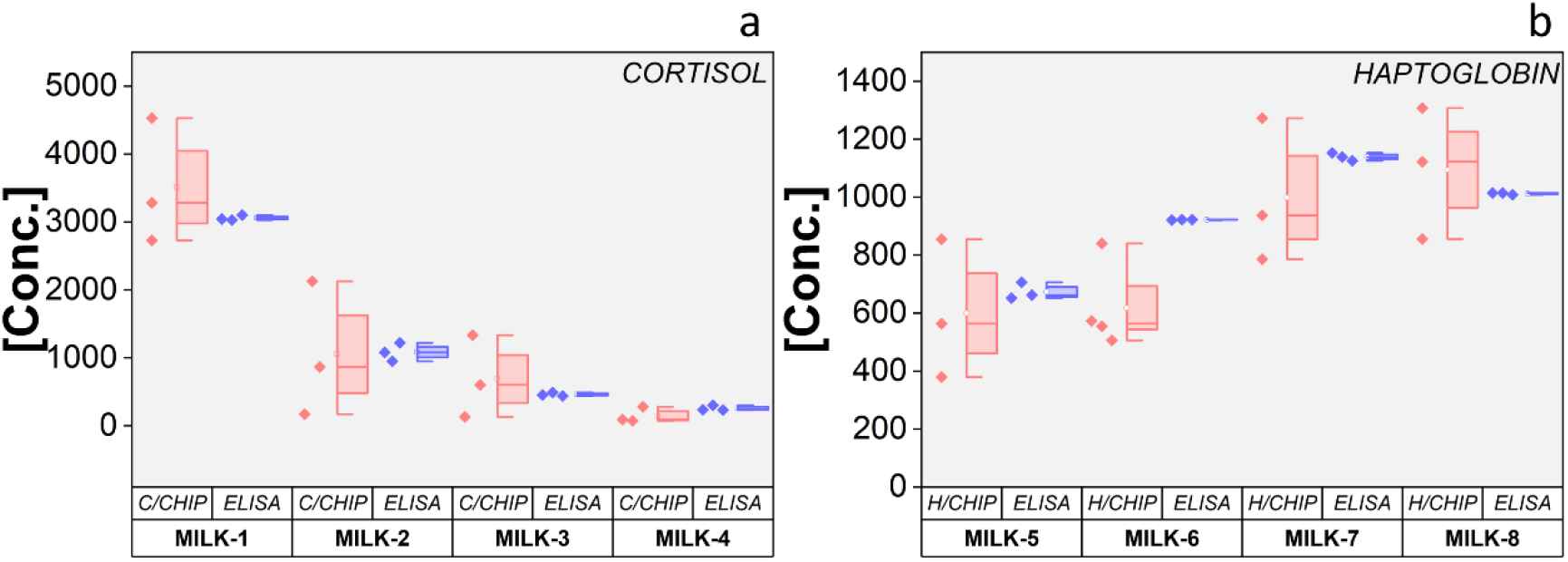
Evaluation of Cortisol and Haptoglobin sensors with milk samples. (a) Cortisol bioassay (C/CHIP) response versus ELISA response using milk samples, (b) Haptoglobin bioassay (H/CHIP) response versus ELISA response using milk samples.

## 4. Conclusion

One of the main advantages of the developed on-chip device is the rapid preparation of the antifouling matrix whose preparation *via* electro-co-deposition takes approximately 2 minutes and the designed device allows achieving electrodeposition of antifouling matrixes onto all sensing electrodes simultaneously. It is possible to integrate this device with a microfluidic system and increase the number of the sensing electrodes for more data acquisition from a single device.

The antifouling coating has abundant functional groups of amines (of chitosan) and carboxyls (of denatured BSA) which makes it suitable matrix for several bioconjugation protocols including carbodiimide chemistry and glutaraldehyde activation. Here we have chosen the high concentration of glutaraldehyde with overnight incubation to achieve high reactivity towards amine groups of protein, however; it is possible to achieve the formation of amide bonding with denatured BSA *via* carbodiimide chemistry (NHS/EDC or sulfo-NHS/EDC) in shorter times such as 30 minutes^30^. Furthermore, lower concentrations of glutaraldehyde (0.5 % glutaraldehyde as shown in Fig. 5b(ii)) provides efficient binding between amine groups of protein and the insoluble chitosan.

In conclusion, we demonstrated that design and fabrication of multiplexed silicon based chips, development of a biocompatible anti-biofouling coating *via* one step electrodeposition and construction of universal electrochemical assay platform for any anti-IgG antibodies. We have shown the broad applicability of the system *via* haptoglobin and cortisol, which are stress biomarkers. The device fabrication is a batch manufacturing process based on our very well established fabrication flows^6-8, 35, 55^. The Aufoam depositions are excellent nanostructures in terms of electrochemistry at ultramicro scale since they increase the surface area due to their high porosity^8^. We have shown the electro-co-deposition of BSA with chitosan in the presence of graphene oxide. This is a straightforward, simple, rapid and reproducible protocol. Such method is an excellent way for selective deposition of materials (nanostructures, hydrogels, and polymers) at micro/nano-scaled surfaces to be used as immobilisation matrices. Developed anti-biofouling coating is effective and provides flexibility for bioconjugation due to abundant functional groups. We have shown the use of amine groups for crosslinking *via* glutaraldehyde chemistry. At this step, we applied a universal recombinant protein of Protein A/G that contains several F_c_ binding domains of both Protein A and Protein G. Therefore, we have established a generic immobilisation platform for a wide range of IgG antibodies. In this study, we have demonstrated the use of the platform with the anti-Haptoglobin and anti-Cortisol antibodies both are IgG isomers. We have shown the performance of the electrochemical bioassays, in buffer and complex matrix. The universal electrochemical bioassay approach can be easily extended to the detection of a wide range of target molecules by simply immobilising the IgG isomer antibodies in the architecture. When coupled with microfluidics, electronic components and wireless communication for data transfer, such system can be applied a panel of target molecules in a complex matrix, leading the way to easy-to-use, rapid and low-cost bioassay platforms.

## Acknowledgements

*This work has been supported by research grants of EU Horizon 2020 (DEMETER 857202), Tyndall National Institute Internal Catalyst Grant (ICA1920), VistaMilk Centre Science Foundation Ireland (SFI) under the grant number 16/RC/3835; Department of Agriculture, Food and the Marine (DAFM) under the grant number 17/RD/US-ROI/56 The authors are also thankful to Dan O’Connell and Anne-Marie Kelleher for their support on the microfabrication of the devices*.

